# A review of operant ethanol self-administration using the sipper model: Methodological advances and a novel standardized analysis tool

**DOI:** 10.1101/2025.08.04.668445

**Authors:** Olivia Ortelli, Jeff Weiner

## Abstract

Despite decades of research, no new FDA-approved medications for alcohol use disorder (AUD) have emerged in over 25 years. Enhancing the translational relevance of preclinical models by more precisely capturing the behavioral and neurobiological features of AUD offers a promising path toward identifying novel therapeutic targets. Operant self-administration paradigms are essential for modeling voluntary ethanol intake in rodents, yet traditional approaches often confound appetitive (seeking) and consummatory (intake) behaviors. The sipper model addresses this limitation by allowing extended, uninterrupted access to ethanol following operant responding, enabling a clearer dissociation between seeking and consumption. In this review, we synthesize key findings from studies employing the sipper model to investigate the behavioral and neurobiological mechanisms underlying alcohol use. We emphasize how over two decades of research employing the sipper model have demonstrated that ethanol- directed behaviors are dynamic processes, shaped by internal states, environmental cues, and prior experience. Finally, we introduce medparser, a new open-source R package designed to standardize the analysis of high-resolution licking data generated by sipper paradigms. By promoting reproducibility and cross-study comparability, this tool supports rigorous behavioral phenotyping. Together, these methodological and analytical advances enhance the translational potential of preclinical models and may ultimately aid in the discovery of novel therapeutic targets for AUD.

## Introduction

Studying alcohol has been of great interest for decades due to the growing understanding of alcohol’s deleterious effects on health and society. Recent statistics indicate that 62% of Americans aged 12 and older consumed alcohol in the past year and 29.5 million individuals (10%) were diagnosed with alcohol use disorder (AUD) (National Survey on Drug Use and Health, 2023). Moreover, alcohol accounted for 3% of all deaths in America during 2020 (99,000 lives) (White et al., 2022). Core symptoms of AUD, as outlined in the 5^th^ edition of the Diagnostic and Statistical Manual of Mental Disorders, include increased craving for or effort spent seeking alcohol, impaired control over alcohol intake, and personal and social impairments as a result of alcohol use (American Psychiatric Association, 2013). Understanding why some individuals develop AUD while others do not, and why some individuals decide to abstain entirely, holds significant clinical and societal importance.

Alcohol self-administration is a complex, multifaceted decision-making process. Understanding the behavioral processes that govern this behavior can yield important insights into the progression from recreational to pathological alcohol use. Preclinical models of alcohol self-administration have delineated three major behavioral components: (1) the reinforcing properties of alcohol, (2) alcohol-seeking (appetitive) behaviors, and (3) alcohol intake (consummatory) behaviors (for review, see Samson & Czachowski, 2003). Importantly, these components are dynamic and modifiable, rather than static.

For instance, the reinforcing value of alcohol depends not only on an individual’s prior history and subjective experience with the drug, but also on the current context. This includes the organism’s physiological state, environmental cues (e.g., presence of alternative reinforcers, effort required to obtain alcohol, and sensory characteristics, like taste), and predictions about the potential consequences of drinking in that moment (Acuff et al., 2023; Acuff, Oddo, et al., 2024; Acuff, Strickland, et al., 2024). Human studies have provided compelling evidence for the role of contextual influences in alcohol-related decision-making (Acuff, Oddo, et al., 2024). While these studies have advanced our understanding of the nuanced and dynamic nature of alcohol use in real-world settings, preclinical rodent models are uniquely positioned to dissect how specific, controlled manipulations affect each facet of the self-administration process. These models also allow for mechanistic investigations of how risk factors (e.g., stress) and protective interventions (e.g., pharmacological treatments) influence reinforcement, seeking, and intake, along with their underlying neural circuitry.

The current review aims to (1) describe the behavioral processes involved with ethanol self-administration, (2) detail preclinical approaches used to study these processes, and (3) review key advances that have emerged from one particularly well-validated methodological approach, the sipper model of operant self-administration. In light of recent developments in data analysis techniques since the introduction of the sipper model, we also highlight how lickometer-based operant paradigms can yield even richer insights. To illustrate this, we present a novel analysis of a compiled dataset generated by our laboratory. Finally, we identify remaining gaps in the literature, with a focus on how preclinical methodologies can be further refined to advance our understanding of ethanol self-administration. Importantly, accurately modeling the core behavioral and neurobiological features of AUD increases the likelihood of identifying promising therapeutic targets.

### 1. Behavioral Processes Controlling Ethanol Self-Administration

Ethanol self-administration is interdependent on two behaviors: 1) it requires one to seek and ultimately obtain ethanol in order to 2) consume ethanol. In fact, appetitive (i.e., seeking) and consummatory behaviors are required for the organization of most behavior, not just ethanol self-administration (Craig, 1918). Appetitive behaviors describe the degree to which an animal directs its behavior in order to procure ethanol (“*how motivated* are you to drink?”, “how *frequently* do you drink?”). These behaviors are shaped by learning and experience, as animals must understand the contingencies which result in ethanol procurement (Samson & Czachowski, 2003). In contrast, consummatory behaviors describe how much and how quickly ethanol is consumed once it’s available. These unconditioned responses are critical for understanding intake control, a core feature disrupted in AUD.

While successful self-administration requires both appetitive and consummatory behaviors, decades of research has demonstrated that appetitive and consummatory processes are differentially influenced by pharmacological and environmental manipulations (see Section 3). Broadly, it is thought that changes in appetitive measures may model changes in wanting or craving while changes in consummatory measures may suggest alterations in post-ingestive, pharmacological effects. While both are essential for modeling ethanol self-administration, dissociating these processes is key to identifying how specific interventions affect motivation versus pharmacological reinforcement.

### 2. Preclinical Models of Ethanol Self-Administration

#### 2.1. Home cage self-administration

Home cage drinking paradigms are among the most used preclinical models of ethanol self-administration due to their simplicity and high throughput. These models include forced and free-choice exposure, delivered on continuous or intermittent schedules (Simms et al., 2008). While forced exposure can overcome neophobia and promote initial intake (Hiller-Sturmhöfel & Kulkosky, 2001; Sardarian et al., 2020), it lacks face validity, as ethanol is the only available fluid. Free-choice paradigms, such as the two-bottle choice test, offer greater translational relevance by allowing voluntary ethanol intake and enabling preference measurements. However, these models do not provide a means to disentangle motivational aspects of alcohol use from consummatory processes.

Although home cage paradigms are methodologically efficient, they offer limited resolution of behavioral processes. Traditional setups often lack precise data on drinking temporal dynamics and lick bout structure, making it difficult to distinguish between changes in seeking versus consumption. Moreover, voluntary intake in home cage models does not reliably predict operant self-administration (Samson & Czachowski, 2003; Wheeler et al., 2025), suggesting distinct underlying behavioral mechanisms. Recent advances, such as the integration of lickometer systems, are beginning to address these limitations by enabling more detailed analysis of drinking behavior with minimal experimenter interference (Godynyuk et al., 2019; Petersen et al., 2023). Importantly, lickometer systems allow for the record of both volume (e.g., number of licks) and temporal dynamics, resulting in the procurement of translationally relevant variables, such as lick rate and drinking microstructures.

#### 2.2. Operant models of self-administration

Operant self-administration procedures are widely used to assess appetitive behaviors alongside consummatory measures. In these paradigms, the completion of a behavioral response (e.g., lever press or nose poke) results in the access of ethanol, typically within an operant box that serves as a discrete self-administration context. Ethanol access is determined by an experimenter-selected schedule of reinforcement. Ratio schedules of reinforcement result in a consequence (i.e., access to ethanol) after a number of behavioral responses are completed while interval schedules of reinforcement result in a consequence following a specific passage of time (Ferster & Skinner, 1957). Under these schedules of reinforcement, the number of reinforcers earned and/or the number of completed behavioral responses are commonly used as the appetitive dependent variable(s). Ethanol delivery methods vary, with the most common still being small-volume dipper presentations, where a motor driven arm raises a cup to deliver a small volume of liquid (∼0.1 mL). This approach is typically paired with a fixed ratio (FR) schedule of reinforcement, resulting in a back-and-forth engagement with the operandum and the dipper. While FR schedules of reinforcement have been widely used to study self-administration of other drug classes, such as stimulants, their application to ethanol self-administration raises unique challenges. The combination of the inherent interruption of drinking with the small volume per delivery may blunt achieved blood ethanol concentrations (BECs), due to BEC being sensitive to both the volume and speed of consumption (Jeanblanc et al., 2019; Samson & Czachowski, 2003). While FR schedules can be effective when paired with sufficient access duration and volume, the use of very small delivery volumes may limit the ability of rodents to achieve meaningful pharmacological effects, particularly within shorter limited access sessions. These limitations highlight a potential divergence between rodent and human self-administration. In humans, alcohol is typically consumed in larger quantities per drinking episode, often in a single sitting without enforced breaks. It is usually ingested for its pharmacological effects, which are experienced more quickly and are paired with more palatable taste components. In contrast, rodent models often assume that animals will expend effort to earn small, often bitter, volumes of ethanol that may not immediately produce reinforcing effects. Such discrepancies may be a key factor in the difficulty of translating preclinical discoveries into clinical interventions.

A further, conceptual limitation of FR operant paradigms is the conflation of appetitive (seeking) and consummatory (drinking) processes. Because ethanol access is contingent on completing a behavioral requirement, it becomes difficult to determine whether a given manipulation, such as a pharmacological treatment or neural circuit intervention, affects motivation to seek ethanol, ethanol consumption, or both. To address these concerns, the “sipper model” was created by Dr. Hank Samson’s laboratory (Samson et al., 1998). Under this paradigm, animals are given 20 minutes to complete a response requirement which results in a retractable, motorized sipper tube to extend into the operant chamber for an additional 20 minutes. For example, a response requirement of 20 is identical to a FR20, where the completion of 20 lever presses results in sipper access. This paradigm is distinct from other FR schedules because the rodent only has the opportunity to complete the FR once, within a fixed time, to obtain “unlimited” ethanol (while there is a finite amount of ethanol in the sipper tube, typically ∼ 35-40 mLs., rodents do not terminate their drinking due to consuming all available fluid, as fluid is remaining in the sipper tube upon termination of the session). The biphasic nature of this model allows for a clear delineation between appetitive and consummatory phases because once the rodent reaches the response requirement, the sipper extends and simultaneously the lever retracts, no longer allowing the rodent to engage with the operandum. This design yields two major benefits when considering ethanol self-administration: 1) appetitive behaviors can be assessed without any influence of ethanol’s pharmacology and 2) the rodent can drink their preferred volume at their preferred speed, without experimenter-induced interruption. Importantly, this design allows for clearer interpretation of behavioral processes and ensures that rodents can achieve pharmacologically relevant BECs within a session.

### 3. Insights from the Sipper Model

The sipper model paradigm has enabled empirical assessment of whether appetitive and consummatory processes involved in ethanol self-administration are functionally independent and/or differentially sensitive to experimental manipulations. Decades of research support these hypotheses (see Table S1). The following sections detail how the sipper model has been employed to dissect the distinct components of ethanol self-administration. Each subsection highlights a specific manipulation or variable that has been instrumental in parsing appetitive and consummatory processes, offering a nuanced understanding of their functional independence and sensitivity to experimental conditions.

#### 3.1. Dissociating appetitive and consummatory processes using the sipper model

A major advancement provided by the sipper model is the ability to study oral ethanol consumption in a way that requires effort to obtain ethanol, while allowing the subject to control the rate and amount consumed without interruption. Unlike paradigms that deliver ethanol in small, discrete volumes (e.g., 0.1 mL per reinforcer), the sipper model supports more naturalistic drinking patterns by minimizing schedule-induced disruptions. For example, Ford et al. (2007) observed shorter latencies to the first lever press and first lick and significantly greater ethanol intake in a self-administration paradigm mirroring the sipper model compared to a traditional FR schedule resulting in 0.1 mL per reinforcer with multiple ethanol presentations.

Beyond its utility in assessing consummatory behaviors, the sipper model has also been used to quantify motivation to seek ethanol without the influence of ethanol’s pharmacological effects, which could confound behavioral responses (e.g,. sedation). Two primary approaches have been developed within this framework to isolate and measure appetitive processes. One approach employs an across-session progressive ratio schedule of reinforcement to derive an index of motivation while maintaining the procedural separation between seeking and drinking (Czachowski & Samson, 1999). In this task, the response requirement to gain access to the sipper tube increases across successive sessions. The primary dependent variable is breakpoint, defined as the highest number of lever presses a rodent completes to earn access. This design allows for daily self-administration sessions in which the animal controls the rate of ethanol intake, yields a reliable motivational metric, and demonstrates strong test–retest reliability after multiple determinations (Czachowski & Samson, 1999). An additional strength of the across-session design is that any failure to meet the response requirement is not confounded by the sedative effects of ethanol or by satiety, since the animal has not yet consumed ethanol in that session. A notable limitation of this approach is the extended time required to determine a single breakpoint value per subject, often spanning several weeks. Additionally, the session in which an animal ultimately fails to meet the response requirement is not known in advance, which limits the feasibility of testing acute pharmacological or environmental manipulations.

To circumvent these limitations, the extinction probe trial (EPT) was developed to quantify a subject’s motivation to procure ethanol and identifies this variable in a single session (Samson et al., 2003). During an EPT, rodents lever press for the entire duration of the appetitive phase (typically 20 min.), yet do not receive ethanol as a consequence of lever pressing. Importantly, there are no cues to signify to the subject that the sipper tube will not extend into the chamber during that session. The total number of lever presses completed within the EPT is the dependent variable used to infer a subject’s motivation to procure ethanol. There is high test-retest reliability when assessing lever pressing during the EPT, allowing for multiple assessments using this behavioral approach (Samson et al., 2001). Moreover, our lab recently demonstrated that the total number of lever presses completed during the EPT is significantly, positively correlated with breakpoints derived from the across session progressive ratio schedule (Ortelli & Weiner, 2024). A striking quality of the EPT that makes it unique compared to other measures of motivation, like breakpoint, is that it operationally separates appetitive and consummatory variables. Many studies have found no relationship between the average intake during the session before or after an EPT and responding during an EPT (Chappell & Weiner, 2008; Czachowski et al., 2003; Samson et al., 2001; Samson et al., 2003, 2004; Samson & Chappell, 2001). Because this measure of motivation can be captured in a single session, the EPT has been used to investigate how pharmacological treatments can mediate motivation to procure ethanol, as well as putative circuits governing motivation to procure ethanol, and are further discussed below (Sections 3.5 – 3.6).

#### 3.2. Sensitivity to response cost (“response requirement”)

Initial studies using the sipper model first probed whether changing the response requirement would affect appetitive and/or consummatory variables. The seminal paper by Samson et al. (1998) was the first to demonstrate that as the response requirement increased, the percentage of rats who met that response requirement would decrease, at the group level. Notably, rats did not adjust their ethanol intake to compensate for the increased effort needed to gain access, suggesting a lack of compensatory drinking in response to greater cost.

These results were replicated in studies modifying this paradigm to implement the across-session progressive ratio schedule (Czachowski et al., 2003; Czachowski & Samson, 1999; Sharpe & Samson, 2003). Serendipitously, in one study Chappell and Weiner (2008) identified a sub-group of rodents (*n* = 6/23) who would not press more than 10-20 times to earn 20 min. access to ethanol. Rather than excluding these rats from the study, the authors allowed these subjects to self-select themselves to a lower response requirement group. These 6 rats were only trained to a response requirement of 8 while the remaining 17 rats were trained to a response requirement of 30. During EPTs, rats trained to the response requirement of 8 exhibited significantly lower levels of lever pressing than those trained to 30, indicating reduced motivation to seek ethanol. Interestingly, average daily ethanol intake did not differ between the two groups, reinforcing the idea that appetitive and consummatory behaviors are dissociable within this paradigm.

Together, these findings underscore the value of using response requirement manipulations to reveal both group-level trends and individual differences in ethanol self-administration. They also suggest that rats engage in context-dependent decision-making processes, whereby environmental constraints (e.g., response cost) influence motivation to seek ethanol without necessarily altering overall consumption.

#### 3.3. Sensitivity to reinforcer solution: Taste, pharmacology, and value

Studies using the sipper model demonstrate that rodents flexibly adjust their behaviors based on the sensory and pharmacological properties of the reinforcer. Comparisons between self-administration behaviors for ethanol, sucrose-sweetened ethanol, and sucrose reveal that both taste and pharmacology contribute to ethanol’s reinforcing value (Carrillo et al., 2008; Czachowski et al., 2003; Czachowski, Prutzman, et al., 2006; Samson et al., 1999, 2000, 2002, 2003; Samson & Chappell, 2002; Sharpe & Samson, 2003), a seemingly obvious yet important detail as humans’ perception of ethanol’s reinforcing strength is also influenced by these reinforced and conditioned stimuli. Indeed, rodents will consume more sweetened ethanol than non-sweetened ethanol (Sharpe & Samson, 2003), highlighting the role of taste on ethanol self-administration. Additionally, ethanol, but not sucrose, preloads can reduce subsequent intake in a dose-dependent manner, indicating behavioral sensitivity to ethanol’s pharmacological effects (Samson et al., 2002).

Within-subject comparisons in this model have also revealed flexible, solution-specific regulation of intake. When rodents were presented with different ethanol concentrations across sessions, they adjusted total volume consumed, particularly during the initial bout of drinking, to maintain a relatively stable g/kg ethanol dose, an effect not observed with sucrose (Samson et al., 1999, 2003; Sharpe & Samson, 2003). These findings collectively suggest that ethanol self-administration in the sipper model is not purely habitual. If lever pressing and drinking were governed by habitual processes, they would be less sensitive to outcome devaluation (Keramati et al., 2011). Outcome devaluation studies, such as pairing ethanol administration with lithium chloride-induced malaise, resulted in significant reductions in ethanol intake and EPT responding the following day (Samson et al., 2004), and therefore are consistent with a goal-directed, value-based decision-making process. Together, these results highlight the utility of the sipper model for dissecting the nuanced, context-dependent mechanisms that govern self-administration.

#### 3.4. Influence of biological and environmental risk factors on self-administration

The sipper model has also been used to investigate how biological and environmental risk factors may influence appetitive and consummatory processes. Many studies have investigated how alcohol preferring (“P”) and high alcohol drinking (HAD1 and HAD2) inbred rat lines behave in this self-administration task (Beckwith & Czachowski, 2014; Bertholomey et al., 2013; Czachowski et al., 2018; Czachowski & Samson, 2002; Henderson-Redmond & Czachowski, 2014; McCane et al., 2018; Verplaetse et al., 2012; Verplaetse & Czachowski, 2015). Studies have demonstrated that P rats have greater appetitive and consummatory behaviors than outbred lines, like Long Evans (Beckwith & Czachowski, 2014; Henderson-Redmond & Czachowski, 2014) and Wistar (McCane et al., 2018) rats. Among all three alcohol preferring lines, P rats have greater appetitive and consummatory behaviors than HAD1 and HAD2 rats, whereas the HAD lines have comparable behavior to one another (Beckwith & Czachowski, 2014; Czachowski & Samson, 2002). These baseline differences highlight the need to consider genetic background when evaluating pharmacological interventions for AUD. Indeed, studies have reported differential effects on appetitive and consummatory behaviors when assessing potential pharmacological therapeutics between P and outbred rats (Henderson-Redmond & Czachowski, 2014; McCane et al., 2018).

In addition to genetic risk, developmental factors like early ethanol exposure can also shape later behavior in this model. Amodeo et al. (2017) reported that rats with a history of home cage ethanol exposure during adolescence had greater appetitive behavior in the sipper paradigm when tested in adulthood compared to rats who did not have a history of ethanol exposure in adolescence. These preclinical findings are congruent with the human literature which states that adolescent alcohol use can have long-lasting consequences, including greater risk of AUD development, which persist throughout adulthood (Spear, 2018). Additionally, McCool and Chappell (2009) reported that male Long Evans rats with a history of social isolation, a commonly used preclinical early life stress model, have increased lever press rates and ethanol intake compared to group-housed control rats. These results may be caused by alterations in stress circuitry, a finding that complements the work of Bertholomey et al. (2013) who reported that yohimbine, a pharmacological stressor, increased reinstatement of ethanol seeking as well as ethanol intake in P and HAD2 male rats. Taken together, combining preclinical models of AUD vulnerability with the sipper model while probing the effects of pharmacological treatments on these populations may ultimately inform personalized treatment approaches for AUD.

#### 3.5. Dissecting pharmacological treatment effects on appetitive and consummatory behaviors

Upon completion of efforts to characterize and validate the sipper model as a preclinical model of oral ethanol self-administration, many studies were conducted to assess how various systemic treatments could influence appetitive and consummatory behaviors. A common finding across these studies was that systemic treatments frequently exert dissociable effects on appetitive versus consummatory behaviors (Czachowski et al., 2001, 2002, 2006, 2018; Ford et al., 2007; Henderson-Redmond & Czachowski, 2014; McCane et al., 2018; Verplaetse et al., 2012; Windisch & Czachowski, 2018).

Additionally, many studies could identify whether treatment effects were specific to ethanol self-administration or whether they generalized to sucrose self-administration (Bach et al., 2023; Butler et al., 2014; Czachowski, 2005; Czachowski et al., 2001, 2012, 2018; Czachowski, Legg, et al., 2006; Czachowski & DeLory, 2009; Ford et al., 2009; Freedland et al., 2001; Henderson & Czachowski, 2012; Henderson-Redmond & Czachowski, 2014; McCane et al., 2018; Sharpe & Samson, 2001; Thorsell et al., 2005; Verplaetse et al., 2012; Verplaetse & Czachowski, 2015; Windisch & Czachowski, 2018). Including sucrose self-administration controls is a key methodological strength of the sipper model. An ideal AUD pharmacotherapy would selectively reduce ethanol seeking or intake while sparing motivation for non-ethanol reinforcers, like sucrose. For example, Czachowski and DeLory (2009) reported that acamprosate reduced ethanol EPT responding in male rats. However, no dose that avoided significant weight loss reduced ethanol intake or affected sucrose-related behavior, suggesting acamprosate may specifically target state-based ethanol craving (Fairbanks et al., 2020). In contrast, naltrexone reduced both appetitive and consummatory responding for ethanol *and sucrose*, indicating a broader, less specific mechanism of action.

#### 3.6. Assessment of brain regions/circuit-specific manipulations

Further studies have used the sipper model to investigate how manipulations of specific brain regions implicated in addiction, such as the nucleus accumbens (NAc), ventral tegmental area (VTA), and amygdala, affect self-administration. Most studies have reported that microinjections of pharmacological agents, typically dopamine antagonists or serotonin agonists, into the NAc (Czachowski, 2005; Samson & Chappell, 2004; Windisch & Czachowski, 2018), VTA (Czachowski et al., 2012), and basolateral amygdala (Butler et al., 2014; McCool et al., 2014) reduced appetitive, but not consummatory, behaviors. These localized interventions allow for greater precision than systemic treatments and support the idea that specific brain regions distinctly modulate different phases of ethanol self-administration.

Even more precise approaches have leveraged modern neuroscience tools to dissect circuit-specific contributions to behavior. For instance, Bach et al. (2023) used chemogenetics to inhibit projections from the basolateral amygdala to the ventral hippocampus and observed reduced EPT responding for both ethanol and sucrose, without altering intake. Similarly, Deal et al. (2020) employed optogenetics to show that tonic stimulation of locus coeruleus norepinephrine neurons increased ethanol intake without affecting seeking, while phasic stimulation decreased both seeking and intake. Budygin et al. (2020) further demonstrated that tonic versus phasic stimulation of VTA to NAc dopamine projections led to opposite effects on EPT responding by decreasing and increasing it, respectively, again without influencing intake. Together, these studies build upon the foundational behavioral work characterizing the sipper model, showing that distinct neural circuits govern appetitive and consummatory phases of ethanol-related behavior.

### 4. MedParser: An Open Source Tool to Analyze Data from the Sipper Model

Recent advances in operant paradigms, particularly the growing use of lickometer-equipped sipper devices, have generated increasingly rich, high-resolution datasets. Despite this influx of detailed behavioral data, there remains a lack of standardization in how these data are processed and analyzed across laboratories. For researchers using lickometers, this lack of standardization can obscure key behavioral metrics, such as microstructural licking patterns or bout dynamics, which are increasingly recognized as translationally relevant. While many laboratories rely on custom scripts to analyze operant self-administration data, a publicly available and customizable R package offers substantial advantages: improving reproducibility, standardizing analytical approaches, and facilitating cross-lab data sharing. To that end, the first author developed medparser, an open-source R package for analyzing data from operant and/or sipper devices (Ortelli & Colarusso, 2024). The package includes a user-friendly Shiny application (available at https://github.com/oortelli/MedParser) which allows users to load raw data, define bout details, extract session-level dependent variables (Table 1), visualize cumulative lick data, and export final datasets to Microsoft Excel for downstream analyses. The application is designed for broad accessibility: users without coding experience can use it via the Shiny interface, while advanced users may modify the source code to tailor it to specific experimental needs.

**Table 1.** Description of Variables Generated to Analyze Self-Administration Behavior from the Sipper Model.

| **Variable** | **Description** |
| --- | --- |
| Latency To First LP | Cumulative time (seconds) elapsed from session start until the first lever press |
| Session Start To Last LP | Cumulative time (seconds) elapsed from session start until the last lever press |
| Time Between First and Last LP | Time (seconds) between the first and last lever press |
| LP Bouts | Number of runs animal was pressing the lever without a 20 second pause between lever pressing<br>*(threshold can be defined by user)* |
| Total LPs | Total number of lever presses within the session |
| LP Rate: Session Start to Last LP | Number of lever presses per minute, time based on the difference between start of session and time of last lever press |
| LP Rate: First to Last LP | Number of lever presses per minute, time based on the difference between the time of the first and last lever presses |
| Latency To First Lick | Cumulative time (seconds) elapsed from session start until first lick |
| Time Between First and Last Lick | Time (seconds) between the first and last lick |
| Total Lick Time | Cumulative time (seconds) rat spends making contact with sipper, excluding time elapsed between drinking bouts |
| Lick Bouts | Number of runs animal was making contact with sipper without a 20 second pause between contacts<br>*(threshold can be defined by user)* |
| Lick Bout Nontrivial | Number of runs animal was making contact with sipper without a 20 second pause between contacts *and* number of events within this bout was greater than 50 *(thresholds can be defined by user)* |
| Total Licks | Total number of licks |
| Lick Rate: First to Last Lick | Number of licks per minute, time based on the difference between the first and last licks |
| Lick Rate: Total Lick Time | Number of licks per minute, time based on total lick time which excludes the time elapsed between drinking bouts |
| First Bout Duration (Percentage) | Percentage of duration (seconds) of first drinking bout compared to total time spent drinking |
| First Bout Lick (Percentage) | Percentage of licks in first drinking bout compared to total number of licks |

*Note.* Abbreviations: LP, lever press

To demonstrate how we can gain new information about consummatory processes by using medparser, 9 days of baseline self-administration sessions from male Long Evans rats responding for 10% ethanol (*n* = 92) or 3% sucrose (*n* = 53) were re-analyzed. All behavioral data were collected by two female research staff members between 2016-2024. For each subject, consummatory and appetitive variables were averaged across completed sessions; sessions in which the response requirement was not met were excluded. However, due to variability in response requirements across cohorts (20, 25, or 30 lever presses), appetitive variables were not included in subsequent analyses to avoid confounding effects. Although prior studies have documented behavioral differences between ethanol and sucrose intake (Section 3.3), no previous work has examined these differences using such a comprehensive dataset. Our aim was to determine which consummatory variables most effectively distinguish ethanol- from sucrose-consuming rats using unsupervised classification methods. To this end, we applied principal component analysis (PCA) and k-means clustering to session-level consummatory data.

Before conducting PCA, we assessed Pearson correlations among variables (Table 1) to reduce redundancy. Lick bouts was removed due to high correlations (|*r|* > 0.80) with three other variables. Similarly, total lick time and percentage of licks completed during the first bout were excluded due to near perfect correlations with total licks and percentage of duration completed during the first bout, respectively. The remaining seven variables (Latency to First Lick, Time Between First and Last Lick, Lick Bouts Nontrivial, Licks, Lick Rate: First to Last Lick, Lick Rate: Total Lick Time, First Bout Duration Percentage) were centered and normalized prior to PCA, which was performed using the R (version 4.3.1) function prcomp(). A visual examination of the scree plot revealed an inflection point after the second principal component (PC), indicating that two components captured the majority of the variance while minimizing overfitting. PC1 explained 49.99% of the variance and primarily reflected overall licking volume, with strong negative loadings for Licks, Lick Bouts Nontrivial, and Time Between First and Last Lick and a strong positive loading for First Bout Duration Percentage. PC2 explained an additional 22.34% of the variance and was driven by variability in licking rate, with high positive loadings for both Lick Rate measures.

To explore whether the PCs captured meaningful group-level structure in the data, k-means clustering (k = 2, seed = 123) was conducted on the retained PCs using the kmeans() function. The choice to use two clusters was supported by the fviz_nbclust() function from the “factoextra” package (Kassambara & Mundt, 2020), which also identified two as the optimal number of clusters. The resulting clusters showed strong alignment with the known group assignments: Cluster 1 included 92.5% (*n* = 49) of the sucrose subjects and no ethanol subjects, while Cluster 2 included all ethanol subjects (*n* = 92) and 7.5% (*n* = 4) of the sucrose subjects (Fig. 1A–B). Silhouette analysis indicated moderate clustering quality for Cluster 1 (silhouette width = 0.65) and strong clustering for Cluster 2 (silhouette width = 0.65), suggesting that ethanol consummatory behaviors were more distinct.

**Figure 1.**
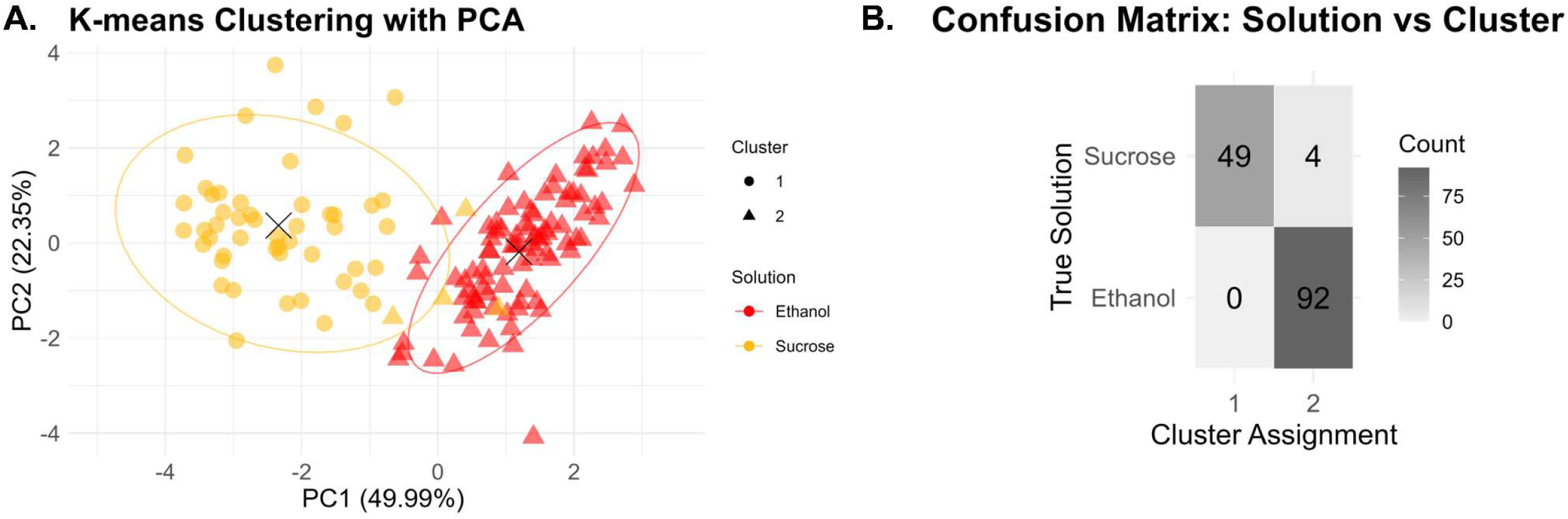
K-means clustering results derived from principle component analysis (PCA). **(A)** Principle components were calculated from session-level consummatory variables of male rats self-administering either 10% ethanol or 3% sucrose. Cluster centers are noted by the corresponding X. Colors represent the actual, known solution the subject was self-administering while cluster assignment from the k-means approach is represented by symbol shape. **(B)** Confusion matrix detailing the number of rats self- administering ethanol or sucrose (“True Solution”) assigned to either cluster from the k-means classification approach (“Cluster Assignment”).

To validate the clustering results, ANOVAs were used to assess whether PC1 or PC2 values were significantly different depending on cluster classification or the known solution the subject was self-administering. PC1 showed significant main effects of both cluster (*F*(1, 142) = 606.03, *p* < .0001) and solution (*F*(1, 142) = 6.40, *p* = .01), indicating that it effectively distinguished both unsupervised and known groupings. PC2 showed a significant effect of cluster (*F*(1, 142) = 6.64, *p* = .01) but not solution (*F*(1, 142) = 1.22, *p* = .27), suggesting that it captured a behavioral phenotype (e.g., binge drinking or front-loading) rather than a solution, per se.

To further character consummatory behavior, we analyzed cumulative lick patterns obtained from the medparser from the final baseline session, focusing on subjects correctly classified by the k-means clustering. Unlike the session-level variables used in PCA, cumulative lick curves offer a temporally resolved view of intake, capturing *how* subjects drink rather than simply *how much*. These curves normalize for total licks and reveal distinct patterns between ethanol and sucrose groups (Fig. 2A). Mann-Whitney tests confirmed significant differences: ethanol rats completed a greater percentage of licks in the first minute (median = 37.92%) compared to sucrose rats (median = 14.08%; *U* = 376, *p* < .0001), and reached 50% of total licks more quickly (median = 1.3 min vs. 5.3 min; *U* = 186.5, *p* < .0001) (Fig. 2B–C). Taken together, these data indicate that ethanol consumption is characterized by early-session drinking, a well-documented phenomenon known as front-loading (see Ardinger et al., 2022 for review). In contrast, sucrose consumption tends to be more evenly distributed across the session, reflecting a steady sipping pattern. Importantly, we have also observed these behavioral patterns in a within-session ethanol vs. sucrose choice paradigm, further supporting that the solutions themselves, rather than prior drinking history, are driving these distinct consummatory patterns (Ortelli & Weiner, 2025). These findings highlight the added value of cumulative lick curves in capturing dynamic features of intake behavior that are not evident from aggregate measures alone.

**Figure 2.**
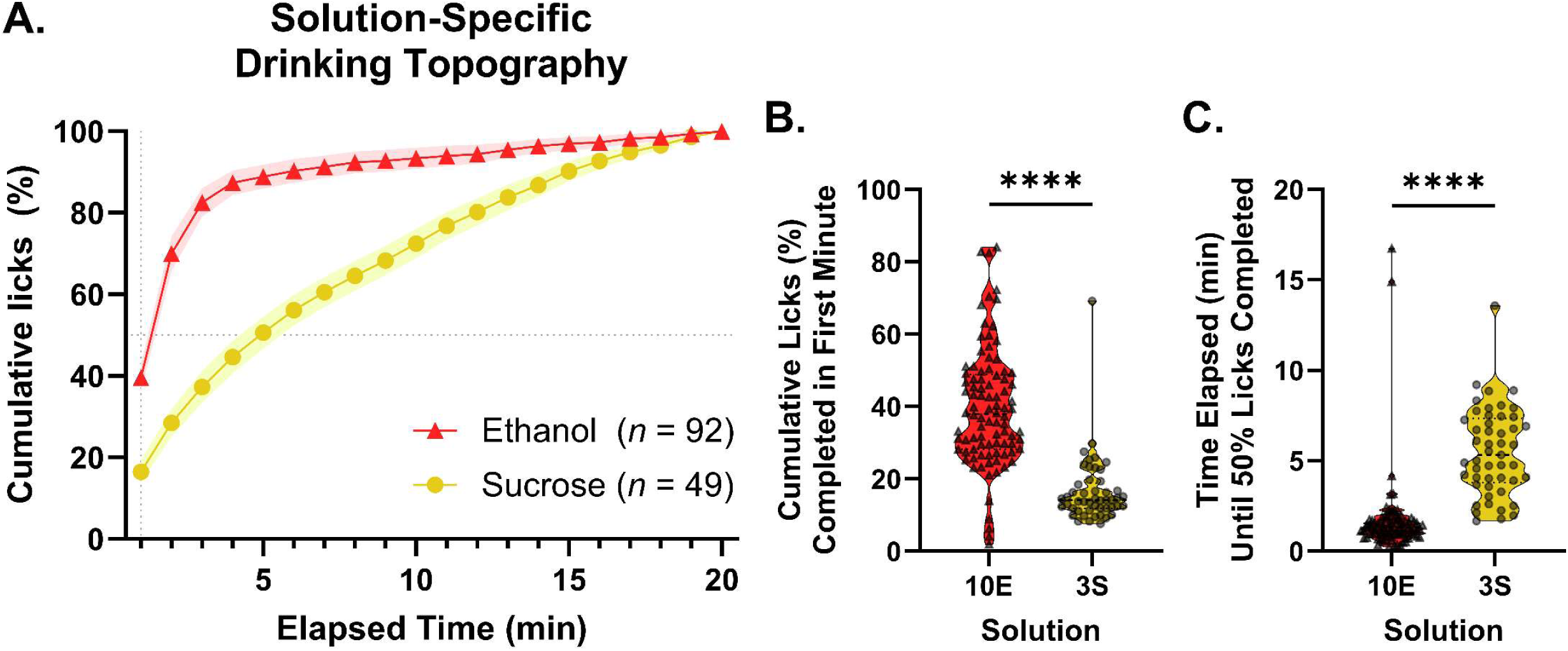
Cumulative lick curve data among the subjects correctly classified from k-means clustering. **(A)** Cumulative lick curve data from a representative baseline session were averaged among correctly- classified subjects self-administering ethanol (*n* = 92/92) or sucrose (*n* = 49/53) (mean with 95% Confidence Interval shading). Subjects self-administering 10% ethanol (10E) and 3% sucrose (3S) demonstrate significant differences in **(B)** the percentage of cumulative licks completed during the first minute of sipper access and **(C)** the time at which half of all drinking was completed. Mann-Whitney tests used to compare distributions of ethanol- and sucrose-drinking rats. \*\*\*\**p* < .0001

While the current data demonstrate the ability to use this code to evaluate consummatory data, this level of analysis can also be completed for appetitive variables. For example, our team recently used an earlier version of this program to demonstrate that inhibiting the projections from the basolateral amygdala to the ventral hippocampus altered lever pressing patterns such that activity in the first five minutes of the EPT, specifically, was especially blunted and never rebounded throughout the duration of the trial (Bach et al., 2023). In addition to being able to analyze lever pressing data from the sipper model, this code was recently adapted to analyze bout behavior of an intravenous self-administration study (Dawes et al., 2024), demonstrating how this code can be extended across other domains of addiction research. Taken together, these analytical approaches offer a powerful and adaptable framework for examining behavioral data, enabling more precise assessments of consummatory and appetitive behaviors across diverse models of addiction research. As open-source code, like the medparser package, continues to be shared across laboratories, they hold the potential to elevate the rigor and reproducibility of behavioral neuroscience research, particularly in the study of substance use.

### 5. Conclusions and Future Directions

In this review, we examined the literature on appetitive and consummatory behaviors in operant ethanol self-administration studies using the sipper model. We also introduced a new open-source analysis tool to quantify these behaviors at higher resolution. Over the past 25 years, our understanding of the environmental and contextual influences, as well as the underlying neurobiological mechanisms, of ethanol self-administration has advanced considerably. However, no new FDA-approved medications for AUD have emerged during this time. These realities underscore the continued need for translationally relevant preclinical models to support the development of more effective AUD treatments. Our analysis tool contributes to the scientific effort to promote standardized, transparent, and reproducible analysis of high-resolution behavioral data. It enables more consistent cross-study comparisons and may help uncover subtle behavioral phenotypes that are critical for identifying novel therapeutic targets.

While the sipper model, and tools like medparser which offer powerful means to quantify nuanced drinking behaviors, have advanced the field, important gaps remain in how these methods have been applied. One notable gap in the literature is the limited investigation into how sex as a biological variable influences appetitive and consummatory phases of ethanol self-administration, and whether the underlying neural circuits are sexually dimorphic. The sipper model is particularly well-suited for investigating sex differences, as it allows subjects 20 minutes of free access to drink at their own pace. This design avoids a key confound present in fixed-volume reinforcement schedules, where administering identical volumes (e.g., 0.1 mL per reinforcer) results in different ethanol doses (g/kg) across sexes due to differences in body weight. Female rats, for instance, require less volume to reach the same g/kg dose as males and would therefore have to lever press less, potentially conflating appetitive and consummatory behaviors.

To date, only two studies have employed both male and female subjects within the sipper model framework (McCane et al., 2018; Ortelli & Weiner, 2024). Our lab’s recent work revealed no sex differences in appetitive responding despite significant differences in intake, a finding likely only possible due to the model’s procedural dissociation of appetitive and consummatory components (Ortelli & Weiner, 2024). McCane et al. (2018) further demonstrated that a candidate pharmacological therapeutic reduced ethanol intake in male but not female P rats, reinforcing the importance of evaluating sex-specific treatment effects. Notably, no studies to date have examined how sex chromosome complement or hormonal mechanisms influence these behaviors, an important future direction that could deepen our understanding of sex as a complex, multidimensional biological construct rather than a binary category (see Grissom et al., 2024).

Another key limitation in current preclinical operant designs is the near-universal use of a single reinforcer, typically either ethanol or sucrose. In these designs, subjects are typically only exposed to one reinforcer, and any comparisons between reinforcers are made across groups rather than within-subjects. This design limits ecological and translational validity, as humans typically have access to alcohol alongside alternative, non-alcohol reinforcers in real-world contexts. Importantly, prioritizing alcohol over alternative reinforcers is a core diagnostic criterion for AUD and other substance use disorders (Banks & Negus, 2017). Future studies, especially those evaluating candidate therapeutics, would benefit from incorporating choice paradigms into operant models to more closely model human decision-making. Our lab has recently developed an operant choice paradigm that leverages the benefits of the sipper model that we believe will be critical in advancing our understanding of the development, maintenance, and modulation of ethanol self-administration (Ortelli & Weiner, 2025). We demonstrate that concurrent ethanol and sucrose availability decreases self-administration of both solutions, particularly ethanol self-administration, and that a manipulation can have solution-specific effects within a single session. This framework could be extended even further to study concurrent self-administration of multiple psychoactive substances, thereby addressing the rising prevalence of polysubstance use, a critical and emerging area of addiction science.

Together, the sipper model and associated analytical tools offer a robust framework for dissecting the behavioral components of ethanol self-administration with greater precision and reproducibility. As the field continues to evolve, integrating these methodological advances with more nuanced experimental designs will be essential for bridging the gap between preclinical research and the development of effective, individualized treatments for AUD.

## Acknowledgements

This work was supported by National Institutes of Health Grants [P50 AA026117, R37 AA17531, R01 AA26551 (JLW), T32 NS115704, F31 AA032154 (OAO)]. All authors have seen the manuscript and approved it for publication. The authors declare that they do not have any conflicts of interest (financial or otherwise) related to the data presented in this manuscript. The authors thank Ann Chappell for thoughtful discussion pertaining to the manuscript. The authors also acknowledge the technical contributions of Ann Chappell and Christina Dyson in collecting the operant self-administration data detailed in this manuscript. Finally, we thank Dr. Hank Samson for his pioneering development of the sipper model and his decades- long contributions to the alcohol research community. His work has profoundly shaped the field of alcohol research and continues to inspire future generations of scientists.

