## Supplemental Table 1 for "A review of operant ethanol self-administration using the sipper model: Methodological advances and a novel standardized analysis tool"

**Table S1**

*Summary of Studies Implementing the Sipper Model to Examine Appetitive and Consummatory Behaviors.*

| Article | Subjects: Strain | Subjects: Sex | Response Requirement | Specific Procedure* | Solution | Manipulation | Change in Appetitive Behaviors | | Change in Consummatory Behaviors | |
|---|---|---|---|---|---|---|---|---|---|---|
| | | | | | | | Ethanol | Sucrose | Ethanol | Sucrose |
| **1** | Long Evans rats | Males | Multiple: 4, 8, 16, 32, 64 | Across-Session Progressive Ratio | 10E or 3S | Response requirement | ↓ | ↓ | – | – |
| **2** | Long Evans rats | Males | Multiple: 4, 8, 12, 16, 20, 25, 30, 40, 50, 60, 70, 80, 90, 100, 120, 140, +20… | Across-Session Progressive Ratio | 10E | Response requirement | | | – | – |
| **3** | Long Evans rats | Males | 4 | Baseline sessions | 10E, 15E, and 20E | Sipper vs. dipper | | | Sipper presentation results in faster rate of intake than dipper | |
| **4** | Long Evans rats | Males | 4 | Extinction Probe Trials | 10E or 3S | | | | First run sizes > Last run sizes | |
| **5** | Long Evans rats | Males | 30 | Extinction Probe Trials | 10E or 3S | Acamprosate treatment | – | – | ↓ | – |
| **6** | Long Evans rats | Males | 30 | Extinction Probe Trials | 10E | Alcohol deprivation effect (2, 7, or 16 days) | – | | – | |
| **7** | Long Evans rats | Males | 10 | Extinction Probe Trials | 10E | Raclopride microinjection into NAc | ↓ | | ↓ | |
| **8** | Long Evans rats | Males | 30 | Extinction Probe Trials | 10E | Repeated extinction probe trials | – | | – | |
| **9** | Long Evans rats | Males | 16 | Baseline sessions | 10E or 3S | Naloxone treatment | ↓ | – | ↓ | ↓ |
| **10** | Long Evans rats | Males | 16 | Baseline sessions | 10E or 3S | SR141716A (selective CB1 antagonist) treatment | ↓ | – | ↓ | ↓ |
| **11** | Long Evans rats | Males | 30 | Extinction/reinstatement trials | 10E or 3S | Self-administered dose (30 second access) of solution following extinction | – | – | | |

| | | | | | | | | | | |
|---|---|---|---|---|---|---|---|---|---|---|
| **12** | P rats, HAD1 rats, HAD2 rats | Males | Multiple: 4, 8, 12, 16, 20, 25, 30, 40, 50, 60, 70, 80, 90, 100, 120, 140, +20…, 600, + 50… | Extinction Probe Trials and Across-Session Progressive Ratio | 10E or 3S | Genetic line | P > HAD1 ~ HAD2 | P > HAD1 > HAD2 | P > HAD1 ~ HAD2 | P > HAD1 ~ HAD2 |
| **13** | Long Evans rats | Males | Multiple: 5, 15, 25, 35, 45, 55, 65, 75, 85, 100, 120, 140, +20… | Across-Session Break Progressive Ratio | 10E or 3S | Concentration manipulation:<br>E: 15, 20, 30, 10 (%)<br>S: 10, 20, 3 (%) | Sweetened Ethanol > Unsweet Ethanol | | Sweetened Ethanol > Unsweet Ethanol | |
| **14** | Long Evans rats | Males | 30 | Extinction/reinstatement trials | 10E or 3S | E group: Preload of 10E or water at varying access times (15, 30, 60, 120s)<br><br>S group: Preload of 3S or water at varying access times (30, 60, 120 s) | ↓ | – | – | – |
| **13** | Long Evans rats | Males | 20 | Baseline sessions | 10E | Nicotine treatment | ↓ | | ↓ | |
| **15** | Long Evans rats | Males | Exp. 1: 20<br>Exp. 2: 4 | Extinction Probe Trials | 10E | Remoxipride treatment | ↓ | | – | |
| **16** | Long Evans rats | Males | 20 | Extinction Probe Trials | Exp. 1: 20E<br>Exp. 2: 10E or 2S | Exp. 1: Multiple EPT sessions | – | – | – | – |
| | | | | | | Exp. 2: Concentration manipulation:<br>→ E group: 0, 2, 5, 10, 15 (%) | – | | – | |
| | | | | | | → S group: 0, 1, 2, 5, 10 (%) | | ↑ | | ↑ |
| | | | | | | → Add 2S to 10E (E group) OR Add 10E to 2S (S group) | – | ↓ | ↑ | ↓ |
| **17** | Long Evans rats | Males | Mutiple: 4, 8, 12, 16, 20, 25, 30, 40, 50, 60, 70, 80, 90, 100, 120, 140, +20… | Across-Session Progressive Ratio | 10E or 3S | | | | E < S | |
| **18** | Long Evans rats | Males | 25 | Extinction Probe Trials | 20E | Devaluation procedure (ethanol gavage + | ↓ | | – | |

| | | | | | | | | | | |
|---|---|---|---|---|---|---|---|---|---|---|
| | | | | | | lithium chloride administration) | | | | |
| **19** | Long Evans rats | Males | 30 | Extinction Probe Trials | 10E | Raclopride microinjection into NAc | ↓ | | – | |
| **20** | Long Evans rats | Males | 4 | Baseline sessions | 10E10S or 10S | | | | Dopamine responses in the NAc are dependent on intake (g/kg), not concentratio n. | |
| **21** | Long Evans rats | Males | 10 (Intake group) or 20 (seeking group) | Intake Assessment or Extinction Probe Trials | 10E or 2S | CGS12066B (Serotonin 1b agonist) microinjection into NAc core | ↓ | – | – | – |
| | | | | | | 8-OH-DPAT (Serotonin 1a agonist) microinjection into NAc core | – | – | ↓ | – |
| **22** | Wistar rats | Males | 20 | Extinction Probe Trials (Fixed Time) | 10E | Chronic intermittent ethanol vapor exposure | ↑ | | ↑ | |
| | | | | | | Neuropeptide Y microinjection into the lateral ventricle | ↓ | | ↓ | |
| **23** | Long Evans rats | Males | 30 | Extinction Probe Trials | 10E or 2S | Baclofen treatment | ↓ | ↓ | ↑ | ↓ |
| **24** | Long Evans rats | Males | 20 | Extinction Probe Trials | 10E | Preload (low vs. high volume, low vs. moderate vs. high dose) | ↓ (dose only) | | ↓ (volume only) | |
| **25** | C57BL/6J mice | Males | 4 and 8 | Extinction Probe Trials | 10E | Schedule of reinforcement | FR schedule with single reinforcer yields shorter response latency than FR schedule with multiple reinforcer presentation s | | FR schedule with single reinforcer yields shorter lick latency and greater intake than FR schedule with multiple reinforcer presentation s | |
| **26** | C57BL/6J mice | Males | 8 | Baseline sessions | 10E | Allopregnanolone treatment | – | | ↑ | |

| | | | | | | | | | | |
|---|---|---|---|---|---|---|---|---|---|---|
| **27** | Long Evans rats | Males | 4 or 20 | Extinction Probe Trials | 10E or 10S | Modified sucrose fade such that rats originally had 3 days of 10S10E exposure then sucrose faded to 10E | | | | |
| **28** | Long Evans rats | Males | 8 or 30 | Extinction Probe Trials | 10E | Baseline sensitivity to ethanol's depressive locomotor effects | – | | ↓ | |
| **29** | Long Evans rats | Males | 30 | Extinction Probe Trials | 10E | Adolescent Social Isolation | Socially isolated animals seek more than group-house controls | – | Socially isolated animals drink more than group-house controls | – |
| **30** | Long Evans rats | Males | 20 | Extinction Probe Trials | 10E or 2S | Acamprosate treatment (No history of chronic intermittent ethanol vapor exposure) | ↓ | – | ↓ (only at high dose; caused health problems) | ↓ (only at high dose; caused health problems) |
| | | | | | | Naltrexone treatment (No history of chronic intermittent ethanol vapor exposure) | ↓ | – | ↓ | ↓ |
| | | | | | | Chronic intermittent ethanol vapor exposure + Acamprosate treatment | ↓ | – | ↓ (only at high dose; caused health problems) | ↓ (only at high dose; caused health problems) |
| | | | | | | Chronic intermittent ethanol vapor exposure + Naltrexone treatment | ↓ | – | ↓ | ↓ |
| **31** | C57BL/6J mice | Males | 16 | Baseline sessions | 10E or 5S | Mecamylamine treatment | ↓ | ↓ | ↓ | ↓ |
| **32** | Long Evans rats | Males | 4 | Baseline sessions | 10E10S or 10S | Solution | | | | |
| **33** | Long Evans rats | Males | 20 (EPT group) or 10 (Intake group) | Extinction Probe Trial or Intake Assessment | 10E or 2S | CNQX (glutamate antagonist) microinjection into the ventral tegmental area | ↓ | – | – | – |
| | | | | | | SCH23390 (Dopamine 1 receptor antagonist) microinjection into the ventral tegmental area | – | – | – | – |
| | | | | | | Tetrodotoxin (voltage-gated sodium channel blocker) microinjection | ↓ | ↓ | – | – |

| | | | | | | | | | | |
|---|---|---|---|---|---|---|---|---|---|---|
| | | | | | | into the ventral tegmental area | | | | |
| **34** | P rats | Males | 20 | Extinction Probe Trials | 10E or 2S | Prazosin treatment | ↓ | ↓ | ↓ | – |
| **35** | Long Evans rats | Males | 10 | Intake Assessment | 10E2S or 2S | Neuropeptide Y microinjection into the central amygdala | – | – | – | – |
| **36** | P and HAD2 rats | Males | 10 | Extinction/reinstatement trials | 10E | Yohimbine (selective alpha-2 adrenergic receptor antagonist) treatment | ↑ | | ↑ | |
| **37** | Long Evans rats | Males | 16 | Extinction Probe Trials | 10E or 3S | BRL (β3-adrenoceptor agonist) microinjection into basolateral amygdala | ↓ | – | – | – |
| **38** | P rats, HAD2 rats, Long Evans rats | Males | 15 | Extinction Probe Trials | 10E | Genetic line | P > HAD2 ~ Long Evans | | P > HAD2 > Long Evans | |
| **39** | P rats, Long Evans rats | Males | 10 | Extinction Probe Trials; Intake Assessment | 10E or 2S | Naltrindole treatment | P rats: ↓ Long Evans: − | P rats: ↓ Long Evans: ↓ | P rats: ↓ Long Evans: − | P rats: − Long Evans: − |
| | | | | | | U50,488H (selective kappa opioid agonist) treatment | P rats: ↓ Long Evans: ↓ | P rats: ↓ Long Evans: ↓ | P rats: ↓ Long Evans: ↓ | P rats: ↓ Long Evans: ↓ |
| | | | | | | Naltrexone treatment | P rats: ↓ Long Evans: ↓ | P rats: ↓ Long Evans: ↓ | P rats: ↓ Long Evans: ↓ | P rats: ↓ Long Evans: ↓ |
| **40** | Long Evans rats | Males | 30 | Extinction Probe Trials | 10E2S | Alpha-m5HT (nonselective serotonin type-2 receptor agonist) microinjection into the basolateral amygdala | ↓ | | – | |
| **41** | Long Evans rats | Males | 4 | Cue-induced reinstatement following 13 days of abstinence | 10E10S or 10S | Age of first ethanol exposure: Adolescents (exposed at post-natal day 36) vs. adults | Adolescents had longer latencies to first lever press and to complete the response requirement but demonstrated no differences in lever pressing | – | – | – |

| | | | | | | | | | | |
|---|---|---|---|---|---|---|---|---|---|---|
| | | | | | | | following forced-abstinence compared to adults | | | |
| **42** | P rats | Males | 20 | Extinction Probe Trials and Intake Assessment | 10E or 1S | Treatment with prazosin, alone or in combination with, propranolol | Prazosin alone: ↓ Propranolol alone: ↓ Prazosin + Propranolol: ↓ | Prazosin alone: ↓ Propranolol alone: ↓ Prazosin + Propranolol: ↓ | Prazosin alone: ↓ Propranolol alone: ↓ Prazosin + Propranolol: ↓ | Prazosin alone: ↓ Propranolol alone: ↓ Prazosin + Propranolol: ↓ |
| | | | | | | Treatment with prazosin, alone or in combination with, naltrexone | Prazosin alone: ↓ Naltrexone alone: ↓ Prazosin + Naltrexone: ↓ | Prazosin alone: ↓ Naltrexone alone: − Prazosin + Naltrexone: − | Prazosin alone: ↓ Naltrexone alone: ↓ Prazosin + Naltrexone: ↓ | Prazosin alone: ↓ Naltrexone alone: − Prazosin + Naltrexone: ↓ |
| **43** | Wistar rats | Males | 16 | Extinction Probe Trials | 20E | Two-bottle choice solutions (water-water vs. 20E-water) and time of exposure (adolescence vs. adulthood) | – | | ↑ | |
| **44** | P rats | Males | 20 | Extinction Probe Trials and Intake Assessment | 10E or 1S | Varenicline treatment | – | – | ↓ | – |
| **45** | Wistar rats, P rats | Males and Females | 10 and 20 | Extinction Probe Trials and Intake Assessment | 10E or 1S | Tolcapone treatment | Male P rats: ↓ Male Wistar rats: ↓ Female P rats: − Female Wistar rats: − | Male P rats: ↓ Male Wistar rats: ↓ Female P rats: − Female Wistar rats: − | Male P rats: ↓ Male Wistar rats: − Female P rats: − Female Wistar rats: − | Male P rats: ↓ Male Wistar rats: − Female P rats: − Female Wistar rats: − |
| **46** | Wistar rats | Males | 10 | Extinction Probe Trials and Intake Assessment | 10E or 2S | Systemic treatment of LY37 (group II metabotropic glutamate agonist) | ↓ | – | – | ↓ |
| | | | | | | Systemic treatment of BINA (selective positive allosteric modulator of group II metabotropic glutamate receptors) | – | – | – | – |

| | | | | | | | | | | |
|---|---|---|---|---|---|---|---|---|---|---|
| | | | | | | Systemic treatment of LY37 + LY37 microinjection into NAc core | ↓ | | – | |
| **47** | Long Evans rats | Males | 30 | Extinction Probe Trials | 10E | Tonic stimulation of ventral tegmental area to NAc dopamine projection | ↓ | | – | |
| | | | | | | Phasic stimulation of ventral tegmental area to NAc dopamine projection | ↑ | | – | |
| **48** | Long Evans rats | Males | 30 | Extinction Probe Trials | 10E | Tonic stimulation of locus coeruleus norepinephrine activity | – | | ↑ | |
| | | | | | | Phasic stimulation of locus coeruleus norepinephrine activity | ↓ | | ↓ | |
| **49** | Long Evans rats | Males | 30 | Extinction Probe Trials | 10E or 3S | Chemogenetic inhibition of basolateral amygdala to ventral hippocampus projection | ↓ | ↓ | – | – |

Abbreviations: E, ethanol; S, sucrose; NAc, nucleus accumbens; P rats, alcohol-preferring rat line; HAD rats, high alcohol drinking rat line. Symbols indicate whether the manipulation increased (↑), decreased (↓), or resulted in no change of (−) the associated behavior.

***Across Session Progressive Ratio:* the response requirement increases across sessions until the response requirement is not met (i.e., breakpoint). *Baseline sessions:* The specified response requirement must be met in order for the sipper to be extended for a fixed time (typically 20 minutes). *Extinction Probe Trial:* regardless of the specified response requirement, a session in which no number of lever presses results in procurement of the sipper tube and the total number of lever presses is recorded. *Intake Assessment:* regardless of the specified response requirement, a session in which only 1 lever press is required for 20 minutes of access to the sipper tube and consummatory behavior is assessed.

**Table S1 References**
